# Draft Genome Sequence of the Asian Pear Scab Pathogen, *Venturia nashicola*

**DOI:** 10.1101/354241

**Authors:** Shakira Johnson, Dan Jones, Amali H. Thrimawithana, Cecilia H. Deng, Joanna K. Bowen, Carl H. Mesarich, Hideo Ishii, Kyungho Won, Vincent G.M. Bus, Kim M. Plummer

## Abstract

*Venturia nashicola,* which causes scab disease of Asian pear, is a host-specific, biotrophic fungus, with a sexual stage that occurs during saprobic growth. *V. nashicola* is endemic to Asia and is regarded as a quarantine threat to Asian pear production outside of this continent. Currently, fungicide applications are routinely used to control scab disease. However, fungicide resistance in *V. nashicola,* as in other fungal pathogens, is an ongoing challenge and alternative control or prevention measures that include, for example, the deployment of durable host resistance, are required. A close relative of *V. nashicola, V. pirina,* causes scab disease of European pear. European pear displays non-host resistance (NHR) to *V. nashicola* and Asian pears are non-hosts of *V. pirina.* It is anticipated that the host specificity of these two fungi is governed by differences in their effector arsenals, with a subset responsible for activating NHR. The *Pyrus-Venturia* pathosystems provide a unique opportunity to dissect the underlying genetics of non-host interactions and to understand coevolution in relation to this potentially more durable form of resistance. Here, we present the first *V. nashicola* draft whole genome sequence (WGS), which is made up of 40,800 scaffolds (totalling 45 Mb) and 11,094 predicted genes. Of these genes, 1,232 are predicted to encode a secreted protein by SignalP, with 273 of these predicted to be effectors by EffectorP. The *V. nashicola* WGS will enable comparison to the WGSs of other *Venturia* spp. to identify effectors that potentially activate NHR in the pear scab pathosystems.

## Introduction

*Venturia nashicola* Tanaka & Yamamoto causes scab disease of Japanese (*Pyrus pyrifolia* var. *culta*) and Chinese (*P. ussuriensis* and *P. bretschneidieri*) pears (Abe *et al.* 2008; Ishii and Yanase 2000; Li *et al.* 2007; Tanaka & Yamamoto 1964). Scab is currently controlled by 15 or more fungicide applications per year (Ishii 1985; Ishii and Yanase 2000). However, resistance to fungicides (such as benzimidazoles and sterol demethylation inhibitors) is an increasing issue (Ishii 2012). There are currently no major commercial cultivars of Asian pear with resistance to *V. nashicola* (Ishii *et al.* 1992; Park *et al.* 2000); therefore, breeding for scab resistance is a high priority for Asian pear breeders. Despite a brief incursion in France (Le Cam *et al.* 2002), *V. nashicola* is limited to Asia, and is regarded as a quarantine threat elsewhere (Jeger *et al.* 2017).

*V. nashicola* (Tanaka and Yamamoto 1964) establishes biotrophic infection of the cuticle and subcuticular space of young leaves and fruit in early spring following rain. Sporulating lesions burst through the cuticle, leading to deformed, cracked, unmarketable fruit and up to 30% yield loss (Lian *et al.* 2006; Lian *et al.* 2007; Eguchi and Yamagishi 2008).

*V. nashicola* was previously regarded as a synonym of *V. pirina* (Aderh.) (synonym, *V. pyrina*) (Sivanesan 1977). Ishii and Yanase (2000) described *V. nashicola* as a distinct species based on morphology, sexual incompatibility with strains of *V. pirina* that infect European pear, and pathogenicity. Species delineation was further supported with molecular analyses (Le Cam *et al.* 2001; Zhao *et al.* 2011). *V. nashicola* and *V. pirina* share a co-evolutionary history with progenitor *Pyrus* hosts, and divergence of the *Pyrus* species has led to differentiation in host specificity in these fungi. *V. nashicola* is restricted to infecting Asian pears, while *V. pirina* is restricted to infecting European pear (Bell 1996; Ishii and Yanase, 2000; Ishii 2002; Abe *et al.* 2008). The incompatible interactions between *V. nashicola* and European pear and conversely, *V. pirina* and Asian pears, are regarded as examples of non-host resistance (NHR) (Heath 2000; Abe *et al.* 2000; Park *et al.* 2000; Bus *et al.* 2013; Won *et al.* 2014). These fungal species are unable to hybridise; however, the host species can form viable hybrids, making the *Pyrus-Venturia* pathosystem excellent for dissecting the role of NHR in determining host-specificity (Won *et al.* 2014). We hypothesize that a subset of the pathogen effector arsenals (secreted proteins that govern virulence and pathogenicity) activate NHR in the *Pyrus-Venturia* pathosystem. Effectoromics has been developed using effector-based screening to identify corresponding resistance (R) receptors in hosts and non-hosts. Effectors of pear scab fungi can be screened to identify and characterize *R* genes that confer resistance to scab fungi (Vleeshouwers *et al.* 2008; Ellis *et al.* 2009). Here we present the first draft whole genome sequence (WGS) of *V. nashicola* (isolate Yasato 2-1-1). The WGS was assembled using mate-paired (MP) and paired-end (PE) reads, with gene annotation assisted by *V. pirina* transcriptome sequences.

## Materials and Methods

### Fungal strain information

*V. nashicola* isolate Yasato 2-1-1, which was sampled from ‘Kousui’ pear (*P. pyrifolia* Nakai) in Ibaraki Prefecture of Japan in 1992 (Ishii *et al.* 2002), was used for genome sequencing in this study. Transcripts from *V. pirina* isolates ICMP 11032 and P35.2 (New Zealand isolates from pear cultivar ‘Winter Nelis’ in Havelock North, Hawke’s Bay, New Zealand) were used to assist *V. nashicola* Yasato 2-1-1 gene prediction.

### Nucleic acid preparation and sequencing

#### DNA preparation and sequencing

Fungal cultures were grown on potato-dextrose agar (PDA) overlaid with a cellophane sheet and on PDA only at 20°C in the dark. Following four weeks of cultivation, cellophane sheets were harvested and mycelia scraped from the sheets, and stored on RNAlater according to manufacturer’s instructions (ThermoFisher Scientific, Rockford,IL). Samples were stored at ambient temperature for shipment from Tsukuba and Minami-awaji, Japan, to Melbourne, Australia. Genomic DNA (gDNA) was extracted from samples using the method described by Kucheryava *et al.* (2008). Quality and quantity of gDNA was determined using a Qubit 2.0 fluorometer (ThermoFisher Scientific) and used to prepare a mate-pair (MP) genomic library using the Illumina Nextera Mate-Pair Protocol (Illumina, San Diego, CA), and a paired-end (PE) genomic library using the KAPA Hyper Prep Kit (Kapa
Biosystems, Wilmington, MA), both with no size selection step. Libraries were sequenced in paired-end mode with 125 bp reads on the Illumina HiSeq1500 at La Trobe University.

#### RNA preparation and sequencing

*V. pirina* conidia were prepared for inoculation as per Chevalier *et al.* (2008). Transcripts generated from *V. pirina* grown in culture were generated under the following conditions: a conidial suspension of ca. 2 × 10^5^ conidia/ml from *V. pirina* isolate ICMP 11032 was used to inoculate PDA overlaid with a cellophane sheet as per Kucheryava *et al.* (2008). Three sheets of inoculated cellophanes per biological replicate were harvested at seven days postinoculation, for a total of three biological replicates. Samples were flash frozen in liquid nitrogen prior to being ground to a fine powder using a mortar and pestle. RNA was extracted following the method described by Chang *et al.* (1993). RNAseq libraries were prepared using an Illumina Truseq stranded mRNA library preparation kit (Illumina) and sequenced in paired-end mode as 125 bp reads on an Illumina HiSeq1500 at La Trobe University. Transcripts from *in planta* were generated as follows: conidia from *V. pirina* isolate P35.2 (isolate used in previous pear scab phenotyping work) were used to inoculate grafted clones of two interspecific hybrids of *Pyrus* (PEAR1 and PEAR2), previously described by Won *et al.* (2014), and two of their progeny (BAG 6305 and BAG 6379), along with *Malus* x *domestica* (cv. ‘Royal Gala’). A conidial suspension of ca. 2 × 10^5^ conidia/ml was applied until run-off to the adaxial side of freshly unfurled leaves with a glass dripper Pasteur pipette, with leaves subsequently enclosed tightly in a plastic snaplock bag for 48 h to enhance the germination of conidia by maintaining a water film. Plants were kept in a glasshouse under ambient conditions with relative humidity (RH) set at 70% and automatic ventilation temperature at 20°C, which resulted in a temperature range of 12-18°C at night and 18-25°C during the day. Leaf samples were collected at three and 10 days post-inoculation, each with three biological replicates. For each biological replicate, five leaves from five separate clonal plants were pooled after harvesting and flash frozen in liquid nitrogen. Frozen leaves were ground to a fine powder in a mortar and pestle under liquid nitrogen. RNA was extracted from 100 mg of ground leaf tissue using the RNA Plant Spectrum Kit (Sigma Aldrich, St. Louis, MI), following the manufacturer’s protocol. RNA extracts of high quality were obtained from all 60 samples with RNA integrity numbers ranging from 5.8 to 8.4 (average of 7.5), as determined using an Agilent RNA chip on a bioanalyzer (Agilent, Santa Clara, CA). RNA from the 60 samples was sequenced at the Australian Genome Research Facility (AGRF) Melbourne on the HiSeq2500 across four lanes of a flow cell.

#### DNA sequence processing methods

Prior to assembly, sequence quality of raw mate pair (MP) reads were checked with FastQC v0.11.2 (Andrews, 2010). PCR duplication level in the sequencing data was assessed and the genome size was estimated using string graph assembler (SGA) v0.10.13/PreQC (Simpson, 2014). The MP data were adapter trimmed using NxTrim v0.4.0 (O’Connell *et al.* 2015) and the reads were separated into multiple categories: the PE reads with short insert size, the MP reads with large inserts, and single reads (SE). The PE sequences were cleaned using Trim Galore v0.4.3 (https://www.bioinformatics.babraham.ac.uk/projects/trim_galore/), where five bases from the 5’-end were trimmed and adapters were culled with a quality cut-off of 28. Overlapping reads in the cleaned PE data were merged using PEAR v0.9.10 (Zhang *et al.* 2014) to build longer reads.

MEGAHIT v1.1.1 (Li *et al.* 2015) was then used to assemble the merged long reads with the rest of the PE data. Furthermore, the resulting contigs were assembled into scaffolds using the MP data with SSPACE-LongRead v2.0 (Boetzer and Pirovano 2014). The genome completeness level was assessed based on the coverage of representative genes in the CEGMA v2.5 (Parra *et al.* 2007) and BUSCO v2.0 (Simão *et al.* 2015) gene sets. RepeatModeler v1.0.8 (Smit *et al.* 2014) was used for the *de novo* identification and classification of repeats in the assembled scaffolds. Thereafter, RepeatMasker v4.0.5 (Smit *et al.* 2015) was used to mask the genome for the newly detected repeats and those in the NCBI repeat database (National Center for Biotechnology Information (NCBI) Resource Coordinators, Database resources of the National Center for Biotechnology Information, 2016).

#### Gene prediction and annotation

Gene expression data from the closely related species *V. pirina,* as well as all Venturiales proteins extracted from the “Identical Protein Groups (IPGs)” (Internet, Bethesda (MD): National Library of Medicine (US), National Center for Biotechnology Information”; 2004, cited 2018 03 01). Available from: https://www.ncbi.nlm.nih.gov/ipg/ were used as supporting evidence.

The *V. pirina* gene expression data were generated from mRNA, derived from two sources: *V. pirina* grown in culture on cellophane sheets, and *V. pirina* grown *in planta* during an infection time-course on interspecific hybrids of *Pyrus* and inoculated *Malus* x *domestica* (cv. Royal Gala). Initially, the *V. pirina* mRNA sequences were mapped to the *V. pirina* genome (Cooke *et al.* 2014) to retrain AUGUSTUS (Stanke *et al.* 2006) and build a parameter profile for *Venturia.* Evaluation of the retraining using the combined set of RNASeq data showed a nucleotide, exon and gene-level sensitivity and specificity of 0.984/0.992, 0.91/0.92, and 0.834/0.857, respectively. 1,834,814 *V. pirina* mRNA reads, and 10,993 Venturiales IPGs were mapped to the *V. nashicola* genome. With these two sets of alignments as hints, and applying the parameter profile trained for *Venturia,* we predicted genes on the repeat-masked *V. nashicola* genome with AUGUSTUS (Stanke *et al.* 2006) via the BRAKER2 pipeline (Hoff *et al.* 2015; Stanke *et al.* 2006).

All annotated predicted proteins were analyzed with SignalP v4.0 (Petersen *et al.* 2011) to predict the presence of N-terminal secretion signals. Predicted secreted proteins were then analyzed with EffectorP v2.0 (Sperschneider *et al.* 2018) to determine the abundance of predicted effector proteins. Gene annotation completeness was assessed using CEGMA (version 2.5) (Parra *et al.* 2007) and BUSCO (version 3.0) (Simão *et al.* 2015) to determine the presence of conserved, single-copy orthologs from the Ascomycota lineage.

#### Data availability

The genome sequencing project has been deposited to the Short Read Archive on NCBI under the accession number SRP144794. The BioProject designation for this project is PRJNA439019.

## Results and Discussion

The draft WGS assembly of *V. nashicola* isolate Yasato 2-1-1 is 45 Mb and consists of 40,800 scaffolds, with a contig N50 value of 69,684 bp (Table 1), resulting in a highly fragmented genome. Despite this high level of fragmentation, BUSCO and CEGMA completeness analyses of the assembled genome revealed 96.9% and 95.97% (respectively) of core fungal genes are represented in this assembly (Table 1).

**Table 1.** *Genome scaffolding statistics*

| Metric | Megahit_scaffolded |
| --- | --- |
| N | 40,800 bp |
| SUM | 45,407,238 bp |
| MIN | 200 bp |
| 1ST-QUARTILE | 231 bp |
| MEDIAN | 231 bp |
| 3RD-QUARTILE | 269 bp |
| MAX | 552,647 bp |
| MEAN | 1112.9225 bp |
| N50 | 69,684 bp |
| CEGMA: |  |
| Complete | 95.97 % |
| Complete + incomplete | 98.79 % |
| BUSCO: |  |
| Complete | 96.9 % |
| Missing | 2.4 % |
| Fragmented | 0.7 % |

There were 11,094 predicted genes (Table 2). Prediction of isoforms was permitted if supported by evidence, but little alternative splicing was predicted, with 10,977 genes with one predicted isoform, 111 with two, and six with three. Overall, 37% of protein-coding gene models are supported with evidence from *V. pirina* protein and/or mRNA evidence along an average of 34% of the length of the transcript.

BUSCO analysis was used to assess the completeness of the predicted gene set. When compared to the “fungi” BUSCO set, 285 of 290 orthologs (98.3 %) were complete and single copy (with one duplicated, three fragmented and one missing). When compared to the “ascomycota” BUSCO set, 1,292 of 1,315 (97.7 %) of orthologs were complete and single-copy (with three duplicated, 21 fragmented and six missing). This shows an improvement of the level of completeness of the total gene set following gene prediction. Therefore, we conclude that this gene set represents an accurate and mostly complete representation of *V. nashicola* genes. The prediction of the secretome resulted in the identification of 1,232 proteins with a putative secretion signal, of these predicted to be secreted proteins, a total of 273 proteins are predicted effectors (Table 2).

**Table 2.** *Gene annotation summary statistics*

| Parameter | Value |
| --- | --- |
| Genes | 11,094 |
| Mean gene length | 1,390.89 bp |
| Max gene length | 26,983 bp |
| Min gene length | 199 bp |
| Mean exons/gene | 2.73 |
| Predicted secreted proteins<sup>a</sup> | 1,232 |
| Predicted effector proteins<sup>b</sup> | 273 |

<sup>a</sup>Proteins harboring predicted signal sequence as determined by SignalP v4.0 (Petersen *et al.* 2011).
<sup>b</sup>Secreted proteins predicted to be effectors as determined by EffectorP v2.0 (Sperschneider *et al.* 2016).

## Conclusions

Here, we present the first draft assembly of the WGS of the Asian pear scab fungus, *V. nashicola*, isolate Yasato 21-1. This draft WGS of *V. nashicola* will enable the identification of candidate effectors and genes involved in fungicide resistance, and contributes to the current genomic resources for *Venturia* spp., specifically *V. pirina* (Cooke *et al.* 2014), *Fusicladium effusum* (pecan scab fungus) (Bock *et al.* 2016), *V. inaequalis* (apple scab fungus) (Deng *et al.* 2017) and *V. carpophila* (peach scab fungus) (Chen *et al.* 2017).

The WGS will enhance comparative genomics between the scab pathogens, in particular with the genome of the closely related European pear scab fungus, *V. pirina* (Cooke *et al.* 2014). These resources will facilitate the identification of effectors involved in host specificity, both within Asian pears, and also the broader NHR of European pear. Effector-assisted breeding will be used to screen for durable disease resistance in pear breeding programs. The WGS will also be useful for identifying essential pathogenicity factors, for example genes governing specific host-pathogen interactions, and the identification of fungicide-targeted protein genes, which will likely be involved in fungicide resistance. Comparative genomics of the pear scab pathogens will also provide an opportunity to identify unique targets for molecular diagnostics of this quarantine pathogen.

## ACKNOWLEDGEMENTS

SJ was supported by a La Trobe University Postgraduate Scholarship, the Plant Biosecurity Cooperative Research Centre, and Japan Student Exchange Scholarships (support from Australasian Plant Pathology Society-the Phytopathological Society of Japan, and the Australia-Japan Foundation, Department of Foreign Affairs and Trade, Australian Government).

DNA sequencing was funded by Kim Plummer, La Trobe University. Assistance and reagents for library preparation for DNA sequencing was provided by Jatinder Kaur, Ross Mann, Fatima Ruma, Josquin Tibbits and Tracie Webster. Sequencing was carried out by the Molecular Genetics Group at Agriculture Victoria in the Department of Economic Development, Jobs, Trade and Resources via La Trobe University Genomics Platform.

Special thanks to Gagan Singla for preparing the grafted plants for the glasshouse work and the Agronomy Team (Yong Tan and Stephen Trolove) and the Post-Harvest Physiology Team (Jason Johnston, Bridie Carr and Matt Punter) for their lab space generosity, and to Brogan McGreal for assessing quality of the RNA samples. RNA sequencing was funded by Plant & Food Research (PFR) Core funding for Pipfruit Research and with the support of “Cooperative Research Program for Agriculture Science & Technology Development (Project No.PJ011913012018)” Rural Development Administration, Republic of Korea.

